# Initial spindle positioning at the oocyte center protects against incorrect kinetochore-microtubule attachment and aneuploidy in mice

**DOI:** 10.1101/2020.06.28.176594

**Authors:** Jessica N. Kincade, Avery Hlavacek, Takashi Akera, Ahmed Z. Balboula

## Abstract

Spindle positioning within the oocyte must be regulated tightly. Following nuclear envelope breakdown (NEBD), the spindle is predominantly assembled at the oocyte center prior to its migration towards the cortex to achieve the highly asymmetric division, a characteristic of female meiosis. The significance of the initial central positioning of the spindle is largely unknown. Here we show that the centered spindle in mouse oocytes is an insurance mechanism to avoid the premature exposure to cortical CDC42 signaling, which perturbs proper kinetochore-microtubule attachments, leading to the formation of aneuploid gametes. Because the spindle forms where NEBD occurs, nucleus position significantly influences the location of the initial spindle assembly. We also find that this nucleus positioning is a dynamic process that depends on maternal age. These findings will help in understanding why female gametes are notoriously associated with high rates of aneuploidy, the leading genetic cause of miscarriage and congenital abnormalities.

## INTRODUCTION

Infertility is a globally prevalent issue affecting approximately 122 million women worldwide (*1*). Errors in chromosome segregation during meiosis result in aneuploidy, the leading genetic cause of miscarriage and congenital birth defects such as Down syndrome (*2, 3*). Aneuploidy occurs in females of all ages, but females of young age (9-20 years) and women of advanced age experience elevated risk of aneuploidy (*4, 5*). Given that chromosome segregation errors occur mostly in female meiosis, particularly during the first meiotic division (Meiosis I, MI) (*6, 7*), it is crucial to understand the molecular mechanisms regulating female MI and how it can go wrong.

Oogenesis is a complex process that begins during fetal development in female mammalian species, shortly after conception. During late pregnancy, oocytes enter MI and duplicate their genome DNA (*8*). Shortly after birth, all germ cells undergo a prolonged arrest at the prophase I stage, which lasts until puberty. In adult ovaries, hormonal cues prior to ovulation stimulate the oocyte to resume meiosis (*8*). The nucleus (germinal vesicle, GV) typically localizes at the oocyte center in fully grown prophase I-arrested oocytes (*9*). Even if the nucleus is off-centered, these cortical nuclei can migrate towards the oocyte center by F-actin-mediated pressure gradient (*10–13*). A recent study suggested that oocytes lacking F-actin with off-centered nucleus positioning have altered transcriptional dynamics (*13*). However, the significance of central nucleus positioning remains largely unknown.

Upon meiotic resumption, prophase I-arrested oocytes undergo nuclear envelope breakdown (NEBD). During NEBD, the microtubule organizing centers (MTOCs) undergo a three-step fragmentation process before their clustering and sorting into two dominant spindle poles (*14–17*). Since the spindle position dictates the cell division plane (*18*), the spindle must migrate to the oocyte cortex to ensure the highly asymmetric meiotic division (*18–22*). Extruding a small polar body (PB) through a highly asymmetric division is critical as it allows the egg to retain the majority of the cytoplasmic storage of maternal RNAs and proteins, which are essential for the early embryonic development (*23, 24*). However, it is still mysterious why the spindle initially forms at the oocyte center, especially since it can increase the risk of symmetric division, leading to infertility.

Here we show that the premature peripheral spindle positioning causes abnormal kinetochore-microtubule (K-MT) attachments, chromosome misalignment and aneuploidy. Importantly, we find that the initial central spindle positioning is critical to avoid the premature exposure to cortical CDC42 signaling, which affects the fidelity of K-MT attachments through microtubule tyrosination. Interestingly, we find that nucleus positioning, a major contributor for spindle location, is a dynamic process that depends on maternal age. The majority of oocytes collected from sexually mature mice had a centrally positioned nucleus, whereas prepubertal and peripubertal mice had more oocytes with a peripherally positioned nucleus, potentially increasing the risk of aneuploidy.

## RESULTS

### Peripheral GV oocytes have higher rates of misaligning chromosomes and forming aneuploid eggs

In mouse oocytes, the meiotic spindle can initially form at the oocyte center (Supplementary Fig. 1A) or near the cortex (Supplementary Fig. 1B) depending on where NEBD occurs. To investigate the impact of the initial spindle position on female meiosis, we sorted fully grown prophase I-arrested oocytes into three groups based on the shortest distance between the nucleus and the cortex: central (larger than 15 µm), intermediate (5 – 15 µm), and peripheral (less than 5 µm) GV oocytes (Fig. 1A,B). Careful rolling of the oocyte, as previously described (*9*), ensured the correct classification. All oocytes selected for the following experiments were comparable in size with a diameter larger than 80 µm (Supplementary Fig. 1C,D), indicating the full-grown status (*25–27*). Any oocyte with a diameter less than 80 µm was removed from further experiments to exclude the possibility of oocyte growth-associated nucleus centration (*9*). DNA staining revealed that all selected oocytes, including peripheral GV oocytes, had a surrounded nucleolus (Supplementary Fig. 1E), a criterion positively associated with developmental competence (*28*). Finally, central GV oocytes were more common in 6-week-old mice compared to intermediate GV oocytes and peripheral GV oocytes (Supplementary Fig. 1F).

**Figure 1:**
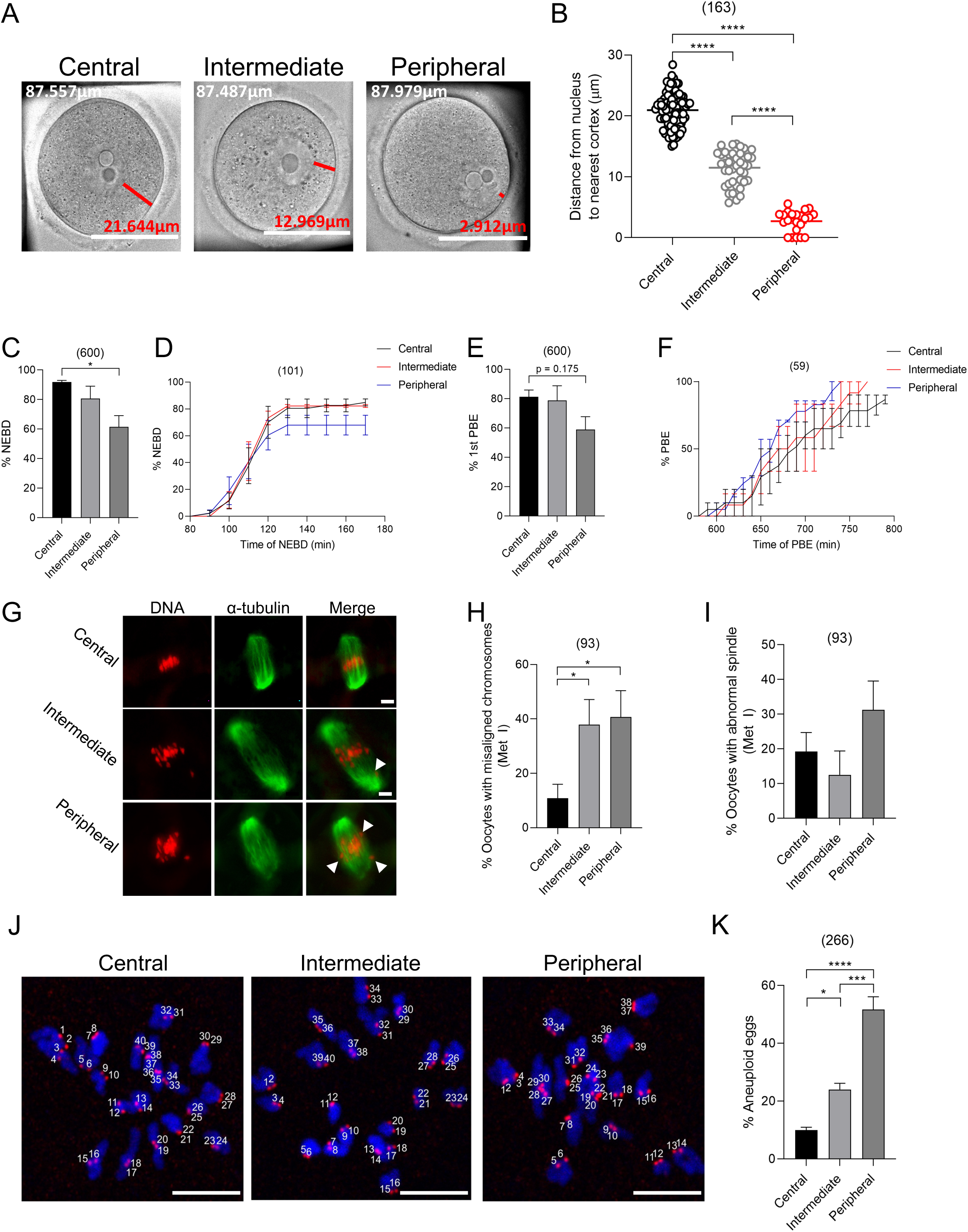
Oocytes with a peripherally located nucleus have higher rates of chromosome misalignment and aneuploidy. (A) Full-grown GV oocytes were collected, imaged, and classified into one of three groups: central GV, intermediate GV, or peripheral GV according to the distance measured from the nucleus to the nearest cortex. The scale bar represents 50 µm. Oocyte diameter is included in the top left of each image in white text; distance from the germinal vesicle to the cortex is included above the scale bar in red text. (B) Quantification of the average distance from the nucleus to the nearest cortex within groups (right graph). (C-F) Full-grown prophase I-arrested oocytes were collected and sorted into central, intermediate and peripheral groups based on nucleus positioning and imaged live using time-lapse microscopy during *in vitro* maturation. (C) Quantification of the percentage of oocytes that underwent NEBD. (D) Quantification of the average time of NEBD. (E) Quantification of the percentage of first polar body extrusion (1st PBE). (F) Quantification of the average time of PBE calculated from oocytes successfully extruded the PB. (G-I) Full-grown prophase I oocytes were collected and sorted into central, intermediate and peripheral groups based on nucleus positioning and *in vitro* matured for 7 h (metaphase I). Oocytes were fixed and immunostained with α-tubulin to label the microtubules (the spindle). DNA was labeled by DAPI. (G) Representative confocal images of metaphase I oocytes. (H) Quantification of the percentage of chromosome misalignment in metaphase I oocytes. (I) Quantification of the percentage of abnormal spindle morphology in metaphase I oocytes. Spindle morphology was assessed using the criterion that a normal meiotic spindle is barrel shaped with two clearly defined spindle poles. (J) Full-grown prophase I oocytes were collected and sorted into central, intermediate and peripheral groups based on nucleus positioning and *in vitro* matured for 14 h (metaphase II). Oocytes were treated with monastrol for 2 h, fixed and immunostained with CREST antibody to label kinetochores. DNA was labeled by DAPI. Oocytes were analyzed individually and scored either as euploid (containing 40 kinetochores) or as aneuploid (containing ± 40 kinetochores). Shown are representative images. (K) Quantification of the percentage of aneuploid eggs. The scale bar represents 10 µm. One-way ANOVA and Tukey’s post hoc tests were performed to determine statistical significance among groups. Data are displayed as mean ± SEM. Values with asterisks vary significantly, * P <0.05, *** P <0.001, **** P <0.0001. The total number of analyzed oocytes (from at least 3 independent replicates) is specified above each graph.

Using the sorted oocytes, we examined the impact of the nucleus position on female meiosis. To examine meiotic progression, oocytes were collected from mice aged 6-8 weeks and matured for 16 h (metaphase II). The majority (∼92%) of central GV oocytes were able to undergo NEBD. In contrast, peripheral GV oocytes showed a significant decrease in NEBD rate (∼61%, Fig. 1C). However, using time-lapse microscopy, we found no significant difference in the timing of NEBD between central GV oocytes and peripheral GV oocytes (Fig. 1D). Similarly, a majority (∼81%) of central GV oocytes were able to extrude the first polar body (PBE), while peripheral GV oocytes were less successful in extruding the first PB (∼60%, Fig. 1E). Once again, there was no significant difference in the timing of PBE between central GV oocytes and peripheral GV oocytes (Fig. 1F). To examine the chromosome alignment and spindle morphology among groups, oocytes were matured *in vitro* for either 7 h (metaphase I) or 16 h (metaphase II). Chromosome alignment was examined based on to parameters previously described (*29*). We found a significant increase in chromosome misalignment at both metaphase I and metaphase II in peripheral GV oocytes, compared to central GV oocytes. Intermediate GV oocytes also showed significantly higher rates of chromosome misalignment when compared to central GV oocytes at metaphase I, but not at metaphase II (Fig. 1G,H and Supplementary Fig. 2A,B). Spindle morphology did not vary significantly among groups either at metaphase I or metaphase II (Fig. 1G,I and Supplementary Fig. 2A,C). Because chromosome misalignment is highly associated with aneuploidy, we then assessed aneuploidy in metaphase II eggs using *in situ* chromosome counting (*30, 31*). Strikingly, segregation errors occurred in peripheral GV oocytes (∼50.15%) approximately five times more frequently than in central GV oocytes (<8 %, Fig. 1J,K). Although we cannot exclude the effect of additional manipulating time (∼ 30 minutes for oocyte sorting/measurements) on oocyte quality, <8% aneuploidy in central GV oocytes remains within the range of aneuploidy in control groups in several previous publications (*32, 33*).

### Incorrect kinetochore-microtubule attachments are increased in oocytes with a peripherally located nucleus

One important cause of aneuploidy is the weakened spindle assembly checkpoint (SAC). Typically, the SAC monitors the status of K-MT attachments and will become inactivated when MTs are correctly attached to all kinetochores. However, a weakened SAC may allow the progression of the cell to anaphase without proper K-MT attachments, leading to aneuploidy. To assess the SAC function in central, intermediate, and peripheral GV oocytes, oocytes were treated with nocodazole, a MT depolymerizing drug that maintains the SAC active and thereby prevents PBE by inducing metaphase I arrest (*31, 34*). We find that central, intermediate, and peripheral GV oocytes all failed to extrude the PB in the presence of nocodazole. In contrast, positive control oocytes treated with ZM447439, an Aurora Kinase B/C inhibitor that is known to disrupt the SAC function at a high dose (10µM), extruded the PB even in the presence of nocodazole (*35*) (Supplementary Fig. 2D). These results suggest that the SAC is intact in peripheral GV oocytes, and therefore, not explaining their observed high aneuploidy rate.

Following spindle migration to the cortex, centromeres facing the cortex have less stable K-MT attachments (*36*). These observations suggest that the cortex can influence MT dynamics and their ability to establish stable attachments with kinetochores (*36*), a critical step to ensure faithful chromosome segregation. In oocytes, even if the SAC is satisfied by present K-MT attachments, it is not necessary that all attachments are correct and homologous chromosomes are bioriented (*29, 37*). To assess K-MT attachments, metaphase I oocytes (∼7 h) were exposed to a brief cold shock (6 minutes) to depolymerize labile non-K-MTs and preferentially maintain stable MT fibers that are attached to kinetochores (*31, 38*). Nucleus positioning had no effect on the percentage of unattached kinetochores (Fig. 2A, Supplementary Fig. 2E). However, peripheral GV oocytes displayed ∼ four-fold increase in the percentage of incorrect K-MT attachments, compared to central GV oocytes (Fig. 2A,B). Moreover, the percentages of peripheral (37.93 ± 9.17%) and intermediate (39.13 ± 10.41%) GV oocytes having two or more abnormally attached kinetochores were significantly higher than those in central GV oocytes (7.14 ± 4.02%, Fig. 2C). These results suggest that improper K-MT attachments are the likely cause, at least in part, of the increased aneuploidy rate in peripheral GV oocytes.

**Figure 2:**
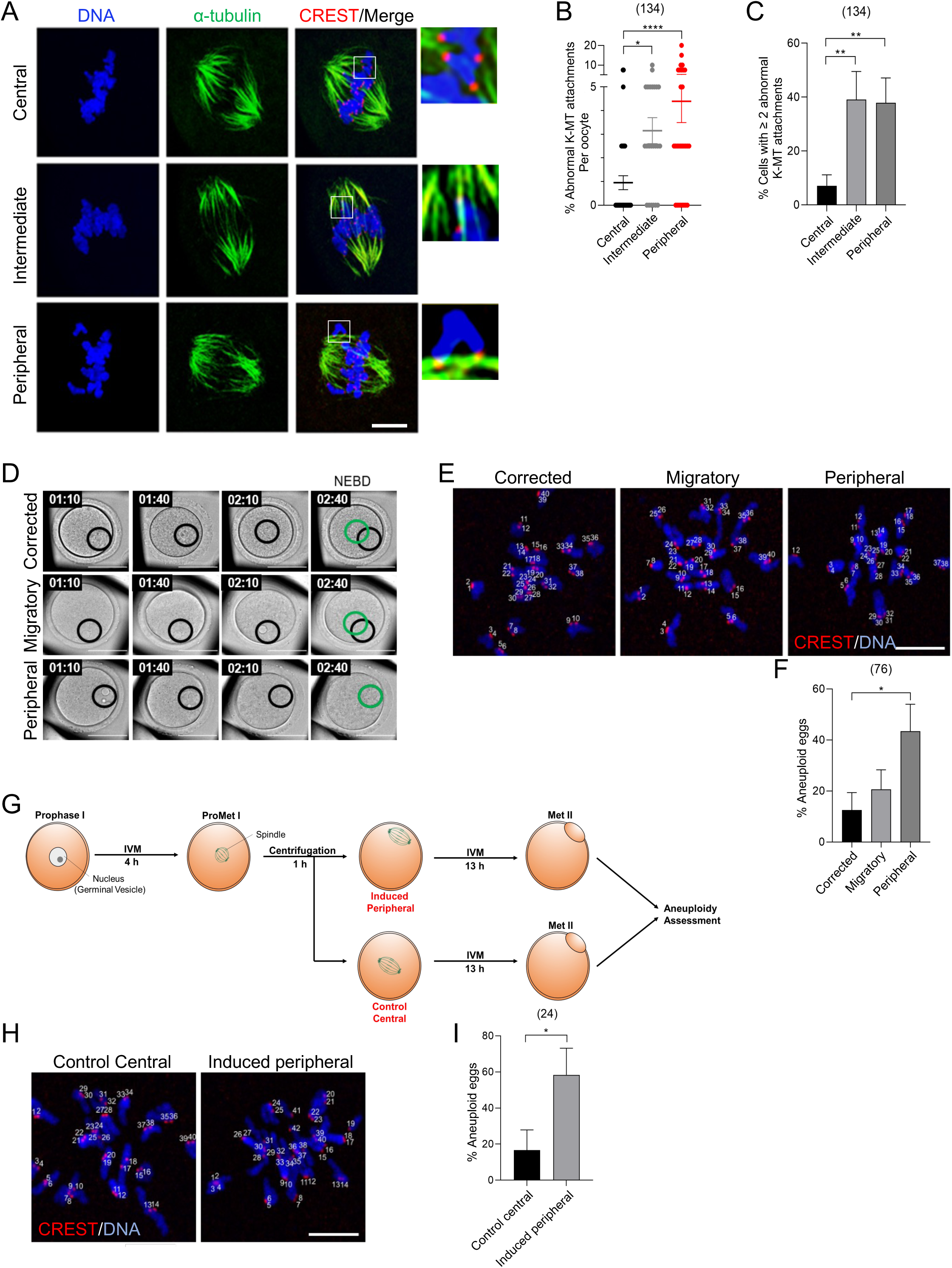
Incidence of incorrect K-MT attachments and aneuploidy associate with spindle proximity to the cortex. (A) Representative images of kinetochore-microtubule (K-MT) attachments in oocytes with a central, intermediate, or peripherally located nucleus matured to metaphase I and immunostained with α-tubulin (to label MTs), CREST (to label kinetochores), stained with DAPI (to label DNA), and imaged using confocal microscopy. Panels on the right (from top to bottom) display normal K-MT attachment, unattached kinetochores and abnormal K-MT attachment. The scale bar represents 10 µm. (B) Quantification of abnormal K-MT attachments. (C) Quantification of the oocytes with 2 or more incorrect K-MT attachments. One-way ANOVA and Tukey’s post hoc test were performed to analyze the data. (D) Representative images of the nucleus movement prior to NEBD. Images were taken every 30 minutes (Z-interval of 5 µm). The black circle denotes the primary nucleus positioning, while the green circle denotes the final nucleus position prior to NEBD. The scale bar represents 50 µm. (E) Representative images of aneuploidy in oocytes experiencing full migration of the nucleus (corrected/central), attempted migration of their nucleus (migratory), or not attempting central migration of the nucleus (peripheral). The scale bar represents 10 µm. (F) Quantification of the incidence of aneuploidy in metaphase II oocytes from the oocytes categorized in D. One-way ANOVA and Tukey’s post hoc test were performed to analyze the data. (G) Schematic depicting the induction of peripheral spindle positioning by mild centrifugation in oocytes collected with a centrally positioned nucleus. (H) Representative images of aneuploidy in oocytes either maintaining central spindle positioning after centrifugation or induced peripheral spindle positioning after centrifugation. The scale bar represents 10 µm. (I) Quantification of the incidence of aneuploidy in oocytes with induced peripherally positioned nucleus and centrally positioned nucleus after centrifugation. Student t-test was performed to determine statistical significance. Data are displayed as mean ± SEM. Values with asterisks vary significantly, * P <0.05, ** P <0.01, **** P <0.0001. The total number of analyzed oocytes (from 3 independent replicates) is specified above each graph.

### Prophase I oocytes favor central nucleus positioning

When meiotic resumption was prevented (by milrinone treatment) in peripheral GV oocytes, the nucleus relocated to the oocyte center within 3 h of incubation in the majority of oocytes (Supplementary Fig. 3A,B and Movie 1), confirming that the nucleus favors the central position, as previously reported (*10, 39, 40*). Central positioning of the nucleus is actin dependent. However, there were no differences observed between oocytes with a centrally or a peripherally located nucleus in terms of cytoplasmic F-actin, cortical F-actin, or perinuclear F-actin (Supplementary Fig. 3C-F). When the oocytes were allowed to resume meiosis, in milrinone-free medium, nuclei initially located at the periphery of the oocyte underwent significant central movement (10 ± 1.256 µm) before undergoing NEBD. Intermediately located nucleus and centrally located nucleus underwent central movement and traveled a distance of 6 ± 0.6024 µm and 4 ± 0.6783 µm, respectively, before NEBD (Supplementary Fig. 4A). However, chromosome alignment in oocytes with a peripherally located nucleus occured relatively closer to the oocyte cortex in comparison to chromosome alignment in oocytes with a central nucleus (Supplementary Fig. 4B,C). It is of note that the timing of chromosome alignment did not vary significantly among groups (Supplementary Fig. 4D), suggesting that peripheral GV oocytes do not exhibit a cell cycle delay. These results suggest that mouse oocytes favor central nucleus positioning, likely to avoid the increased incidence of aneuploidy in peripheral GV oocytes.

### Incidence of aneuploidy directly associates with the proximity of early spindle positioning to the cortex

Because the spindle forms where NEBD occurs, whether at the center (Supplementary Fig. 1A) or at the cortex (Supplementary Fig. 1B), it is likely that increased aneuploidy in peripheral GV oocytes is, at least in part, due to peripheral spindle formation. Using live time-lapse microscopy, we found that oocytes with a peripherally located nucleus demonstrate one of three types of movement: oocytes that were able to achieve complete relocation of the nucleus to the center prior to NEBD (the spindle is formed centrally) were termed ‘corrected’, oocytes that demonstrated central movement prior to NEBD but did not achieve full relocation were termed ‘migratory’, and oocytes that failed to make a relocation attempt before NEBD and remained at the cortex were termed ‘peripheral’ (the spindle is formed peripherally; Fig. 2D). Strikingly, although all oocytes selected in this experiment were peripheral GV oocytes, oocytes that were able to successfully relocate their nucleus to the center and experienced central spindle formation showed relatively lower rates of aneuploidy when compared to oocytes that failed nucleus relocation and underwent NEBD at the periphery of the oocyte (Fig. 2E,F). These results suggest that peripheral GV oocytes are not inherently erroneous, but instead peripheral spindle formation is detrimental to the fidelity of chromosome segregation.

After meiotic resumption, mouse oocyte spindles are typically formed and remained at the oocyte center until late metaphase I (∼7h) followed by their timely migration towards the cortex. It is technically feasible to shift the position of the spindle within the oocyte without noticeable adverse effects (*22, 41*). To further confirm the relationship between the aneuploidy rate and the proximity of early spindle positioning to the cortex, we experimentally relocated the spindle prematurely to the cortex through mild centrifugation (Fig. 2G), a technique proven successful in murine oocytes (*42*). This centrifugation technique has been used extensively for several purposes (*e.g.,* lipid droplet removal and prior to parthenogenetic activation) without noticeable detrimental effects on oocyte developmental competence (*43–45*). Central GV oocytes were selected and matured *in vitro* for 4 h, until prometaphase I (the time prior to establishing correct K-MT attachments) (*46*). These prometaphase I oocytes were centrifuged, examined live under the microscope and resorted into two groups according to their spindle positioning: those maintaining central positioning (control central) and those with induced peripheral positioning (induced peripheral), which was followed by *in vitro* maturation for an additional 13 h to reach metaphase II. Importantly, although all oocytes were originally central GV oocytes, oocytes with induced peripheral positioning showed significantly higher rates of aneuploidy when compared to oocytes maintaining central spindle positioning after centrifugation (Fig. 2H,I), indicating that the premature cortical proximity of the spindle (prior to establishing correct K-MT attachments) causes erroneous chromosome segregation.

### Premature exposure to the cortical CDC42 signal perturbs K-MT attachments in oocytes

A conserved Rho-family GTPase, CDC42, is the master regulator of cell polarity in multiple systems. To investigate whether the premature exposure of the spindle to the cortical cues perturbs the establishment of correct K-MT attachments, we inhibited the cortical CDC42 signaling in prometaphase I oocytes with a peripherally located spindle. Peripheral GV oocytes were matured *in vitro* for 5 h, and oocytes with a peripherally positioned spindle (confirmed using Z-series live imaging) were selected and treated with ML141 (a previously proven selective CDC42 inhibitor in mammalian oocytes) (*47–49*) for an additional 2 h (Fig. 3A). Examining K-MT attachments in metaphase I oocytes showed that the number of unattached kinetochores did not differ significantly among groups (Supplementary Fig. 5A). Inhibition of CDC42 by using ML141 has no significant effect on cytoplasmic or cortical F-actin in oocytes with a peripherally located spindle (Supplementary Fig. 6A-C) nor on K-MT attachments in oocytes with a centrally located spindle (Fig. 3B,C). Importantly, inhibition of cortical CDC42 signals using ML141 partially rescued abnormal K-MT attachments in oocytes with a peripherally positioned spindle (Fig. 3B,C). To further confirm this finding, we perturbed the CDC42 signaling by overexpressing CDC42 dominant-negative mutant (CDC42 T17N, previously demonstrated to inhibit CDC42 in mouse oocytes) (*50*). Consistent with the ML141 results, perturbing CDC42 signaling by overexpressing a CDC42 dominant-negative mutant significantly rescued abnormal K-MT attachments in oocytes with a peripherally located spindle (Fig. 3D,E). Taken together, these results suggest that the increased incidence of abnormal K-MT attachments in oocytes with a peripherally located spindle is, at least in part, due to the premature exposure to cortical CDC42 signaling.

**Figure 3:**
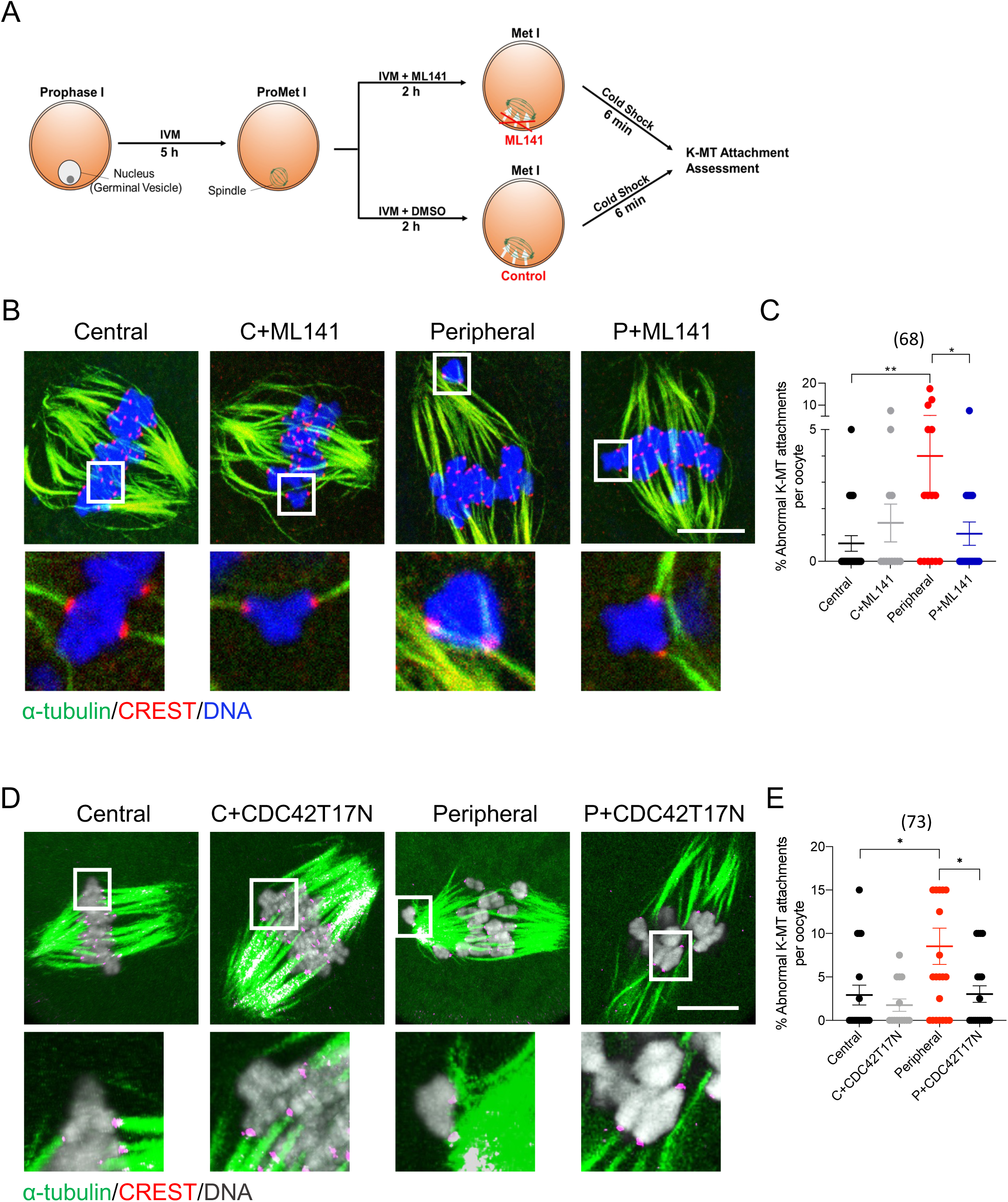
Cortical CDC42 signaling impairs proper K-MT attachments in oocytes with peripherally located spindles. (A) Schematic depicting the treatment of collected oocytes with ML141, a CDC42 inhibitor, in order to inhibit the cortical influence upon kinetochore-microtubule (K-MT) attachment formation. (B) Representative images of DMSO and ML141-treated oocytes matured to metaphase I and immunostained with α-tubulin (to label MTs), CREST (to label kinetochores), stained with DAPI (to label DNA), and imaged using confocal microscopy to assess K-MT attachments. (C) Quantification of the average incidence of abnormal K-MT attachment in B. (D) Representative images of K-MT attachment in oocytes expressing CDC42 dominant negative mutant (CDC42 T17N). The scale bar represents 10 µm. (E) Quantification of the average incidence of abnormal K-MT attachment in D. The scale bar represents 10 µm. One-way ANOVA and Tukey’s post hoc test were performed to determine statistical significance. Data are displayed as mean ± SEM. Values with asterisks vary significantly, * P <0.05, ** P <0.01. The total number of analyzed oocytes (from 3 independent replicates) is specified above each graph.

Induced peripheral oocytes (*i.e.* oocytes with a peripherally located spindle after centrifugation) exhibited higher rates of abnormal K-MT attachments, compared to those that maintained a centrally positioned spindle (control central) after centrifugation (Supplementary Fig. 5B,D). To confirm that K-MT attachments in induced peripheral oocytes were affected primarily by the proximity to the cortex and to exclude the possibility that the difference in the response to centrifugation between induced peripheral and control central oocytes was due to a pre-existing difference, we asked whether inhibiting CDC42 cortical signals could rescue the increased incidence of abnormal K-MT attachments in induced peripheral oocytes. Control central and induced peripheral prometaphase I (5 h) oocytes were treated with or without ML141 for an additional 2 h prior to assessing K-MT attachments (Supplementary Fig. 5B). There were no differences in the percentage of unattached kinetochores among groups (Supplementary Fig. 5C). Importantly, CDC42 inhibition largely rescued the increased incidence of abnormal K-MT attachments in induced peripheral oocytes (Supplementary Fig. 5D), further supporting the idea that increased abnormal K-MT attachments in induced peripheral oocytes is due to the premature exposure of the spindle to the cortical CDC42 signal. Collectively, these results strongly suggest that avoiding the cortical CDC42 signal during prometaphase I (*i.e.* by being at the oocyte center prior to timely spindle migration) is critical in protecting the oocyte’s ability to establish correct K-MT attachments, leading to faithful chromosome segregation.

### Excessive MT tyrosination interferes with establishing correct K-MT attachments

We next asked how CDC42 affects K-MT attachments. Because the cortical CDC42 plays an essential role in actin cap formation by activating ARP2/3 complex (*51*), we tested whether CDC42 signaling perturbs K-MT attachments in peripheral GV oocytes via the premature activation of ARP2/3. To address this question, peripherally GV oocytes were matured *in vitro* for 5 h and oocytes with a peripherally positioned spindle (confirmed using Z-series live imaging) were treated with CK-666 (a selective and potent ARP2/3 inhibitor) (*51, 52*) for an additional 2 h. Analyzing K-MT attachments in metaphase I oocytes showed that ARP2/3 inhibition did not rescue abnormal K-MT attachments in oocytes with a peripherally located spindle (Supplementary Fig. 6D,E), indicating that CDC42 signaling does not perturb the establishment of K-MT attachments via actin cap formation.

Recent study using mouse oocytes suggested that CDC42 signaling can promote α-tubulin tyrosination within the cortical spindle, increasing MT dynamics (*36*). Therefore, we hypothesized that the premature exposure of the spindle to cortical CDC42 (*i.e.* before establishing correct K-MT attachments) results in excessive MT tyrosination, which interferes with the correction mechanism of K-MT attachments. The genetically encoded C-terminal tyrosine on α-tubulin undergoes cycles of detyrosination and (re) tyrosination (*53*). In mitotic cells, initial incorrect attachments are established during early mitosis and primarily formed by tyrosinated MTs. Tyrosinated MTs are relatively labile to depolymerization, leading to their detachment from chromosomes. When correct attachments are established, MTs are detyrosinated and stabilized (*54*). In mouse oocytes, reducing α-tubulin tyrosination (by depleting tubulin-tyrosine ligase) increased the rate of MT stabilization (*36*). Thus, the level of MT tyrosination must be spatiotemporally regulated to establish correct K-MT attachments. To test whether α-tubulin tyrosination levels depend on the spindle position, we compared tyrosinated α-tubulin levels in metaphase I oocytes between central and peripheral spindles. In contrast to β-tubulin levels, which were not affected by the spindle position, tyrosinated α-tubulin levels were significantly higher in oocytes with a peripherally positioned spindle when compared to oocytes with a centrally positioned spindle (Fig. 4A,B). We also confirmed the impact of CDC42 signaling on α-tubulin tyrosination in the spindle. Our results showed that tyrosinated α-tubulin levels were significantly lower in CDC42-inhibited oocytes (oocytes expressing CDC42 dominant negative mutant, CDC42 T17N), compared to control oocytes (Fig. 4C,D). Thus, CDC42 cortical signaling promotes MT tyrosination in mouse oocytes. We then asked whether decreasing tyrosination levels could rescue increased abnormal K-MT attachments in peripheral GV oocytes. Removing the tyrosine residue from α-tubulin (*i.e.* detyrosination) is regulated by tubulin carboxypeptidase enzymes encoded by Vasohibins (VASHs) (*55, 56*). Ectopic expression of VASH2 did not completely abolish tyrosinated α-tubulin in mouse oocytes (Fig. 4E), whereas co-expression of VASH2 and small VASH binding protein (SVBP, a chaperone protein required for VASH stability) (*55, 56*) eliminated tyrosinated α-tubulin without affecting the bipolar spindle formation (Fig. 4E-G, Supplementary Fig. 7). Importantly, decreasing α-tubulin tyrosination significantly decreased abnormal K-MT attachments in oocytes with a peripherally located spindle (Fig. 4H,I). Taken together, our data indicate that the premature exposure of the spindle to cortical CDC42 signaling interferes with the establishment of correct stable K-MT attachments, via promoting MT tyrosination.

**Figure 4:**
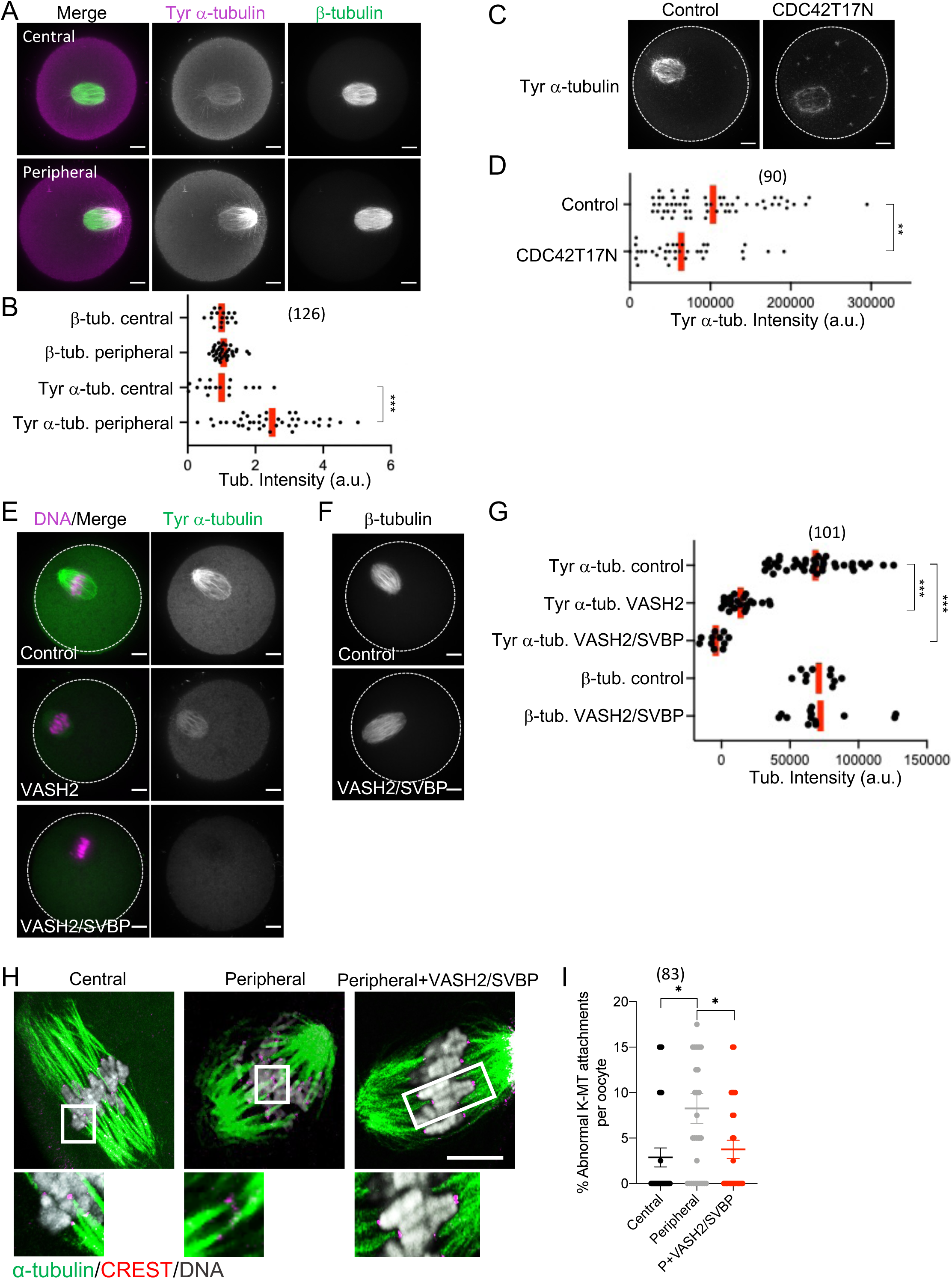
Reducing tubulin tyrosination in peripheral GV oocytes rescued the abnormal K-MT attachments. (A) Representative images of oocytes with centrally and peripherally located spindles stained for tyrosinated α-tubulin and β-tubulin. (B) Quantification of tubulin intensities within the spindle. (C) Representative images of oocytes expressing CDC42 dominant negative mutant (CDC42 T17N) stained for tyrosinated α-tubulin. (D) Quantification of tyrosinated α-tubulin levels within the spindle. (E,F) Representative images of oocytes expressing VASH2 and VASH2/SVBP stained for tyrosinated α-tubulin (E) and β-tubulin (F). (G) Quantification of tubulin intensities within the spindle. (H) Representative images of K-MT attachment in peripheral GV oocytes expressing VASH2 and VASH2/SVBP. (I) Quantification of abnormal K-MT attachments in H. The scale bar represents 10 µm. One-way ANOVA and Tukey’s post hoc test were performed to determine statistical significance. Data are displayed as mean ± SEM. Values with asterisks vary significantly, * P <0.05, ** P <0.01, *** P <0.001. The total number of analyzed oocytes is specified above each graph.

### Too young female mice tend to have peripheral GV oocytes increasing the risk of aneuploidy

It is established that the egg aneuploidy rate increases with advanced maternal age mainly due to chromosome cohesion loss. However, it is becoming increasingly clear that too young females also have higher chances of producing aneuploidy eggs(*5*), though the underlying mechanisms are largely unknown. We wondered if the nucleus position may explain the higher aneuploidy rate in younger females. To test this idea, we quantified the nucleus position *in vivo* by analyzing ovary sections from mice with different ages (2-, 3-, 5-, and 8-week-old) (Fig. 5A-C). We found that the majority of oocytes were categorized as central GV oocytes for the 8-week-old mice, but the younger mice (2-, 3-, and 5-week-old) had fewer central GV oocytes, and the peripheral GV oocytes were the majority in the 2- and 3-week-old mice (Fig. 5A-C). *In vitro*-cultured oocytes from 2-, 3-, 5-, and 8-week-old mice showed similar trends with the *in vivo* experiment (Fig. 5D). These results suggest that oocytes from very young mice tend to have a peripheral nucleus, possibly increasing their aneuploidy risk through the premature exposure of the spindle to cortical cues.

**Figure 5:**
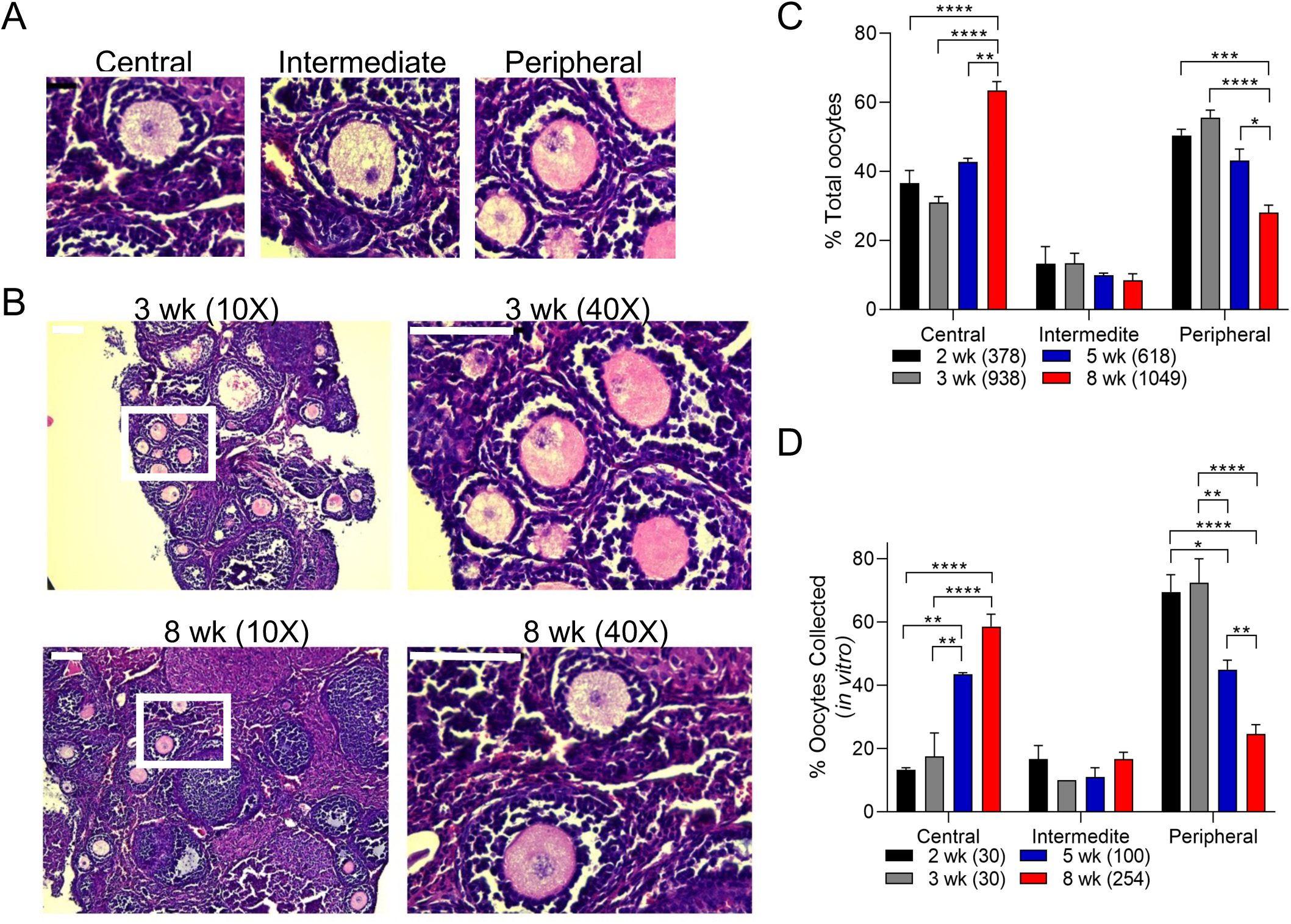
Nucleus positioning is a dynamic process throughout the development. (A) Histological analysis of nucleus positioning *in vivo* at different ages. Ovaries from 2, 3, 5, and 8 weeks old mice were collected, fixed, and stained with Hematoxylin and Eosin. Sections were observed through 10X and 40X objective magnification and classified into one of the three positioning groups: central, intermediate, peripheral. The scale bar represents 100 µm. (B) Representative images of nucleus positioning in 3 and 8 weeks old mice. The white box emphasizes the location of the magnified image within the ovary. The scale bar represents 100 µm. (C) Quantification of the *in vivo* proportion of oocytes sorted according to their nucleus positioning (central, intermediate, peripheral) in 2, 3, 5, and 8 week old mice. (D) Quantification of the in vitro proportion of oocytes sorted according to their nucleus positioning (central, intermediate, peripheral) in 2, 3, 5, and 8 week old mice. Two-way ANOVA and Sidak’s multiple comparison test were performed to determine if the groups differed significantly. Data are displayed as mean ± SEM. Values with asterisks vary significantly, * P <0.05, ** P <0.01, *** P <0.001, **** P <0.0001. The total number of analyzed oocytes (from 3 independent replicates) is specified above each graph.

## DISCUSSION

Aneuploidy occurs frequently in female oocytes, typically as a result of mis-segregation during MI (*6, 7, 33, 57*). We developed different complementary and confirmatory oocyte models in which: 1) the spindle is assembled peripherally or centrally (based on the nucleus position or the location of the spindle at NEBD), 2) the centrally positioned spindle is prematurely relocated towards the cortex (using centrifugation), or 3) cortical CDC42 signaling is perturbed in oocytes with a peripherally located spindle. We found that initial spindle positioning at the oocyte center (until proper K-MT attachments are established and prior to timely spindle migration) is an insurance mechanism to avoid the premature exposure to cortical CDC42 signaling, which hinders proper K-MT attachments, leading to aneuploidy. These findings may explain why, in mouse oocytes, correct K-MT attachments are primarily established when the spindle is still at the oocyte center (∼ 2 h before anaphase I), *i.e.* before migrating to the cortex (*36, 46*).

Nucleus positioning in meiotic oocytes is often studied only *in vitro* or at a single time point *in vivo* and, thus, the comprehensive understanding across different timepoints is missing. Also, there has been a long-standing debate regarding the position of the nucleus in mouse oocytes. For example, a previous study found that the majority of prophase I-arrested oocytes have a peripherally located nucleus, and this cortical positioning of the nucleus is likely influenced by gap junctions and increased cumulus cell contact (*58*). On the other hand, other studies demonstrated that the majority of prophase I-arrested oocytes have a centrally positioned nucleus (*9, 13, 59*). Interestingly, in the first study (*58*), the oocytes were collected from mice aged 21 days, whereas in the other studies (*9, 13, 59*), the oocytes were collected from mice aged more than 6-8-weeks old. Our results clearly link these previous findings, providing an explanation to the long-standing debate on the nucleus position in mouse oocytes and advancing our understanding of an important phenomenon.

Although the nucleus shifts from a peripheral position in prepubertal oocytes towards a central position in sexually mature mouse oocytes, this is likely not the only shift during the reproductive life. Indeed, it has been shown that oocytes from aged mice (∼1 year) also exhibit a majority of peripherally positioned nuclei (*9*). These observations indicate that oocyte nucleus migrates to the oocyte center during sexual maturation and migrates back to the cortex with the advancement of maternal age. Investigating the molecular basis underlying the age-dependent nucleus migration would be an interesting future direction. For example, analyzing cytoskeleton dynamics at different developmental stages may reveal why the nucleus migrates towards certain directions at certain stages.

Nucleus relocation within eukaryotic cells is critical for organism development. In *Drosophila* oocytes, the nucleus migrates from a central posterior to an asymmetrical anterior position to dictate the dorsal-ventral axis, necessary for organogenesis and *Drosophila* development (*60–62*). In *C. elegans* zygotes, the two pronuclei meet at the posterior cortex and then migrate towards the zygote center to allow proper spindle pole orientation along the anterior-posterior axis prior to spindle migration towards the posterior cortex (*63–66*). Although many mammalian oocytes at prophase I are not polarized and the nucleus position has no consequence on the developmental axis, they still employ a similar nucleus relocation. The peripheral nucleus undergoes centration which correlates with maturation success. Following NEBD, the centrally formed spindle must migrate towards the cortex in a timely manner to ensure the asymmetric division. Our understanding of the biological significance of central nucleus positioning in mammalian oocytes is still rudimentary. It has been proposed in Almonacid *et al*. (2018) that nucleus centration plays a role in keeping prophase I oocytes in a non-polarized state, ensuring random cytoplasmic distribution and avoiding unphysiological cytoplasmic compartmentalization (*11*). Our findings suggest that central nucleus positioning is an insurance mechanism to allow the avoidance of the spindle to the premature exposure to cortical cues, providing a complementary explanation of why mouse oocytes experience nucleus centration.

Current *in vitro* fertilization (IVF) technologies have much room for improvement, as only 12.7% of naturally ovulated IVF cycles in humans lead to pregnancy and only 8.8% of those pregnancies result in live births (*67*). Although technologies and protocols have advanced since the first IVF birth in 1978, the IVF success rate remains less than 50% for women under the age of 35 (*68*). To date, there are no definitive morphological markers of predictive oocyte aneuploidy. Gametes from human females are notoriously associated with high rates of aneuploidy (*4, 6*). In addition, human oocytes with a peripherally located nucleus are associated with poor maturation rates compared to those with a centrally located nucleus (*59*). Therefore, it is tempting to speculate that central nucleus positioning is a strategy to ensure initial spindle positioning at the oocyte center, thereby avoiding the development of aneuploid gametes. Understanding the association between peripherally located spindle and the incidence of aneuploidy may allow us to better screen for the oocytes predisposed to proper chromosome segregation.

## MATERIAL AND METHODS

### Ethics

All laboratory animals were managed, and experiments conducted, in compliance with the University of Missouri (Animal Care Quality Assurance Reference Number, 9695).

### Mouse strains

Unless otherwise specified, female CF-1 WT mice were used. *Cep192-eGfp* reporter mice were generated (*69*) by integrating a construct harboring the EGFP reporter gene into CF-1 mouse genome via CRISPR/Cas9-mediated homology-directed repair. The EGFP reporter was fused at the C-terminus of the endogenous mouse Cep192. Homozygous breeding pairs of *Cep192-eGfp* reporter mice were used to maintain the colony.

### Oocyte collection, microinjection and in vitro maturation (IVM)

Full-grown GV oocytes were collected from CF-1 female mice aged 6-8 weeks. Oocyte collection and culture were carried out as previously described (*31, 70*). Cumulus oocyte complexes (COCs) were collected and cultured in bicarbonate-free minimal essential medium (MEM) supplemented with 3mg/ml polyvinylpyrrolidone (PVP) and 25 mM HEPES (pH 7.3) under mineral oil (MilliporeSigma, St. Louis, MO, USA # P2307, # H3784, # M8410). Oocytes were manually denuded and sorted into 3 groups: Central GV, Intermediate GV, Peripheral GV oocytes. Careful rolling of the oocyte, as previously described, ensured correct classification (*9*). Denuded GV oocytes were microinjected with ∼5 pl of the indicated cRNA using Eppendorf Femtojet 4i. The injection medium for oocytes was MEM supplemented with 3mg/ml polyvinylpyrrolidone (PVP), 25 mM HEPES (pH 7.3) under mineral oil. Oocytes were then transferred to Chatot, Ziomek, and Bavister (CZB) maturation medium (*71*) supplemented with 1µM glutamine (MilliporeSigma, # G8549) under mineral oil and matured at 37°C with humidified 5% CO_2_ in an incubator for either 7, 14, or 16 h.

Nocodazole (Sigma #M1404), monastrol (Sigma #M8515), ML141 (Sigma #SML0407), ZM447439 (Tocris #2458) and CK-666 (Sigma #182515) were dissolved in DMSO and added to CZB culture medium at a final concentration of 5 µm, 100 µM, 500 nM, 10 µm and 50 μM, respectively. SiR-tubulin and SiR-DNA were added to the maturation medium (at a final concentration of 100 nM) during live imaging to label MTs and DNA, respectively.

### In vitro cRNA synthesis

Synthesis of cRNAs was performed using T7 polymerase (mMessage mMachine kit, Ambion) according to the manufacturer’s instructions. The cRNA was purified using an RNAEasy kit (Qiagen). cRNAs used for microinjection were 3xFLAG-Vash2 (mouse VASH2 with three tandem FLAG tag at the N terminus) at 500 ng/μl, SVBP-HA (mouse SVBP with a HA tag at the C terminus) at 300 ng/μl and *Cdc42 T17N* (dominant-negative CDC42 mutant) at 800 ng/μl.

### Immunocytochemistry

After maturation, oocytes were fixed in a freshly prepared solution of 3.7% paraformaldehyde (MilliporeSigma, # P6148) dissolved in phosphate buffer saline (PBS). Oocytes were incubated in permeabilization solution (0.1% Triton X-100 in PBS) for 20 minutes and blocking solution (0.3% BSA and 0.01% Tween-20 in PBS) for 20 minutes before staining. Oocytes were incubated at room temperature for one h with the primary antibody before undergoing 3 consecutive washes in blocking solution for 9 minutes each. Oocytes were then incubated at room temperature for one h in the secondary antibody solution. Phalloidin (ThermoFisher Scientific #T7471; 1:50) was added to the secondary antibody solution to label F-actin.

Following another 3 consecutive washes in blocking solution at 9 minutes each, oocytes were mounted on glass slides using Vectashield containing 4’,6-Diamidino-2-Phenylindole, Dihydrochloride (DAPI; Vector Laboratories, Burlingame, CA, USA) in order to stain the DNA. Fluorescence was observed using a 40X oil objective using Leica TCS SP8 confocal microscope or Leica DMI8 fluorescence microscope and oocytes were imaged using 3 µm Z-intervals. Oocytes were analyzed using NIH ImageJ software (National Institute of Health, Bethesda, MD USA).

The following primary antibodies were used in immunofluorescence: conjugated α-tubulin-AlexaFluor 488 (Life Technologies # 322 588; 1:75), CREST autoimmune serum (Antibodies Incorporated # 15-234; 1:25), tyrosinated α-tubulin (BIO-RAD# MCA77G; 1:500), and anti β-tubulin (9F3) monoclonal conjugated to Alexa Fluor 488 (Cell Signaling Technology, #3623; 1:50).

### In situ chromosome counting

Oocytes matured *in vitro* for 14 h in CZB medium supplemented with L-glutamine under mineral oil were transferred to CZB containing 100 µm monastrol (MilliporeSigma, # M8515) and incubated an extra 2 h; monastrol is an Eg5-kinesin inhibitor which induces monopolar spindle formation and thus results in a rosette of chromosome distribution (*30, 31*). After treatment with monastrol, oocytes at metaphase II stage were fixed in a freshly prepared solution of 2.5% paraformaldehyde (MilliporeSigma, # P6148) and stained with CREST autoimmune serum (Antibodies Incorporated # 15-234; 1:25) to label the kinetochores. DAPI was used to label the DNA. Fluorescence was observed using a 40X oil objective using a Leica DMI8 microscope and oocytes were imaged using 0.5 µm Z-intervals in order to observe all kinetochores. Oocytes were analyzed individually and were scored either as euploid (containing 40 kinetochores) or as aneuploid (containing ± 40 kinetochores).

### Assessment of kinetochore-microtubule attachment

Cells were matured *in vitro* for 7 h in CZB medium supplemented with L-glutamine under mineral oil. The oocytes were placed on ice for 6 minutes in a 96 well dish containing chilled MEM. Exposing the oocytes to cold shock depolymerizes unattached MTs; however, when MTs have established end-on attachment to a kinetochore, it becomes cold stable (*38*). Oocytes were fixed using a freshly prepared 2.5% paraformaldehyde solution and immunostained with anti-human CREST (to label kinetochores) and α-tubulin (to label MTs) antibodies. DAPI was used to label DNA. Cells were imaged using a Leica TCS SP8 confocal microscope and a x63 oil objective, taking images at 0.5 µm Z-intervals. Kinetochores were scored as normal (attached to bioriented chromosomes), unattached, or abnormally attached (syntelic - kinetochores of both homologous chromosomes attached to one side of the spindle or merotelic - a kinetochore maintaining attachment to both sides of the spindle), as previously described (*31*). Oocytes were analyzed using NIH ImageJ Software.

### Time-lapse microscopy

Full-grown oocytes were imaged using a 40X oil objective on Leica TCS SP8 confocal microscope or Leica DMI8 microscope within a microenvironmental chamber maintaining an atmosphere of 5% CO_2_ and temperature of 37°C in humidified air. Imaging acquisition began before NEBD. Images were captured at 5 µm Z-intervals. Image sequences were analyzed using NIH ImageJ Software.

### Ovary collection and histological evaluation

Ovaries from CF-1 mice of varying ages (2 weeks, 3 weeks, 5 weeks, and 8 weeks) were collected, submerged, and stored in buffered zinc formalin overnight. Tissues were then moved to 70% ethanol for storage until undergoing a 10-h protocol using a Leica ASP300S enclosed tissue processor. After processing, tissues were embedded in paraffin wax and cooled fully using the Leica EG1150 C/H pair. Embedded tissues were sectioned at 8 µm thickness using a Leica Histocore AutoCut microtome. Representative samples were taken for each age by taking 3 consecutive sections periodically (every 4, 6, 8, or 10 sections respectively). Tissues were stained with Hematoxylin and Eosin (Abcam, # 245880) before being mounted with permount resin (Thermo Fisher Scientific, Inc. # SP15). Oocytes and their corresponding nucleus positioning were examined using 10X or 40X objective on a Leica DMI8 microscope.

### Statistical analysis

Tests used to evaluate the statistical significance of the findings reported include One-way ANOVA and Student t-test using GraphPad Prism. The Tukey post hoc test was utilized to determine statistical significance between groups. Student’s t-test was performed for normally distributed numerical data following D’Agnostino-Pearson omnibus normality test. P values of < 0.05 were considered significant. All data are displayed as means ± SEM.

### Declaration of interest

All authors declare that there are no conflicts of interest that may prejudice or bias the research reported.

## Funding

This research was supported by laboratory start-up funding from the University of Missouri and the NIH (R35 GM142537) to AZB and the Division of Intramural Research at the National Institutes of Health/National Heart, Lung, and Blood Institute (1ZIAHL006249-01) to TA.

## Acknowledgements

The authors would like to thank all members of the Balboula lab at the University of Missouri, Columbia for dedicated help and encouragement. The authors would like to thank Rocio Rivera, Heide Schatten for critical reading of the manuscript and valuable discussion, Yuksel Agca, Pramod Dhakal, Thomas Spencer, the Animal Modeling Core and the Molecular Cytology Core for valuable discussion and equipment usage, and Guillaume Halet for providing the CDC42 T17N construct.

## Author Contributions

AZB conceived the project. AZB, JK and TA designed and analyzed experiments. JK, AH, TA and AZB performed experiments. AZB and JK wrote the manuscript. All authors read/edited the manuscript.

## Supplementary Figures

**Supplementary Figure 1:**
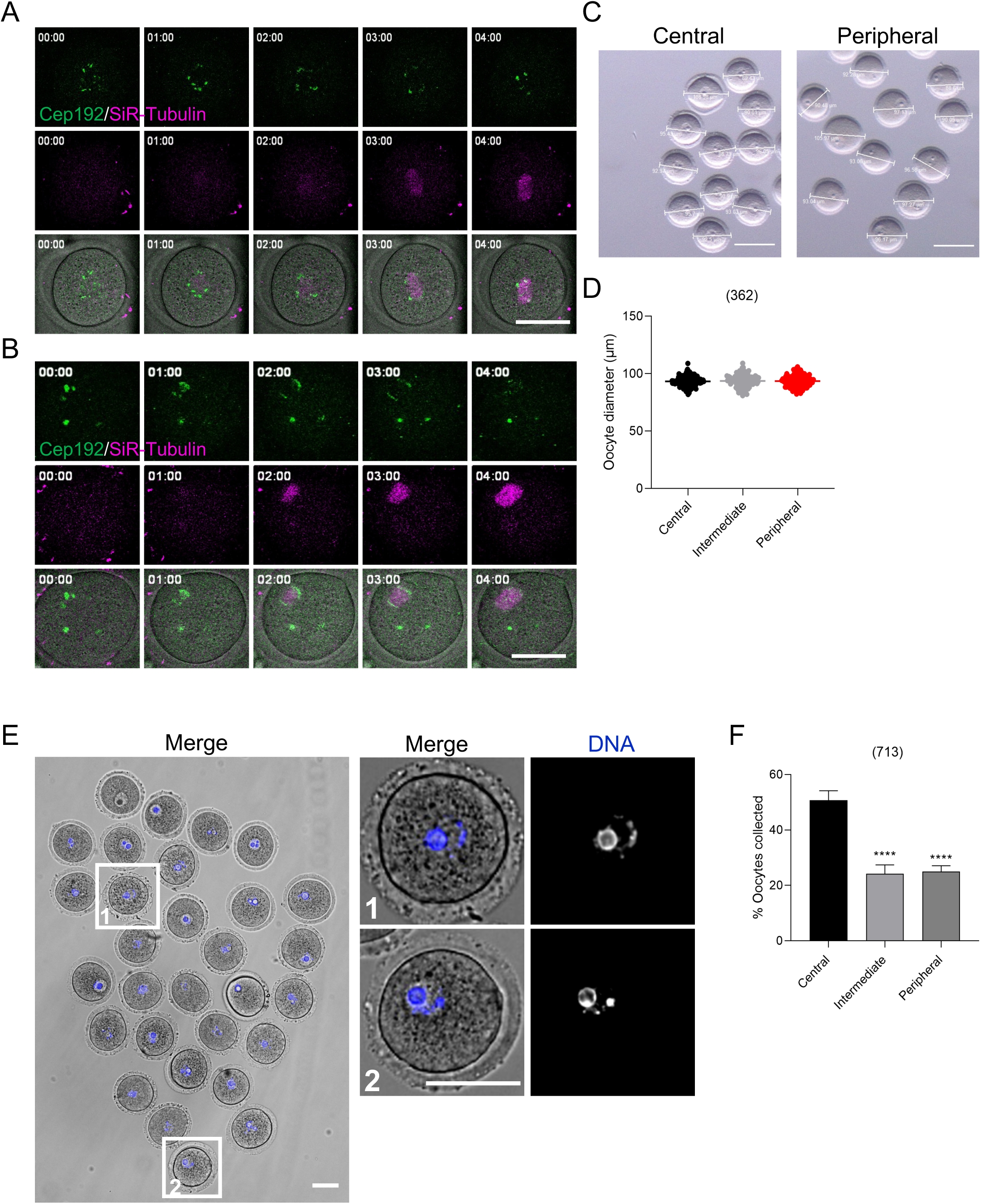
Full-grown prophase I-arrested oocytes were collected from *Cep192-eGfp* reporter mice, sorted into central (A) and peripheral (B) groups based on the positioning of their nucleus and *in vitro* matured in CZB medium containing SiR-tubulin (to label microtubules, magenta). Spindle poles were labeled by CEP192-EGFP (MT organizing centers, green). Oocytes were imaged live using time-lapse confocal microscopy. Shown are representative fluorescence and bright-field (lower panels) image sequence depicting the location of NEBD and consequent spindle formation. Images were taken every 1 h (Z-projection of 13 sections every 5 μm). The scale bar represents 50 μm. (C) Oocytes were sorted based on location of their nucleus into central or peripheral groups, imaged and quantified according to their diameter to ensure similarity between groups. Please note that the large oocyte (105.97 μm, peripheral group) was undergoing cell death and excluded from further experiments. (D) Quantification of the average diameter of oocytes collected for experimentation. (E) Representative fluorescent image of Hoechst staining (to label DNA) in group of selected, but not sorted, oocytes. Representative images of oocytes with a peripherally located nucleus and a surrounded nucleolus are featured in the right panel. The scale bar represents 50 μm. (F) Quantification of nucleus position in oocytes. One-way ANOVA and Tukey’s post hoc test were performed to analyze the data. Data are displayed as mean ± SEM. Values with asterisks vary significantly, **** P <0.0001. The total number of analyzed oocytes is specified above each graph.

**Supplementary Figure 2:**
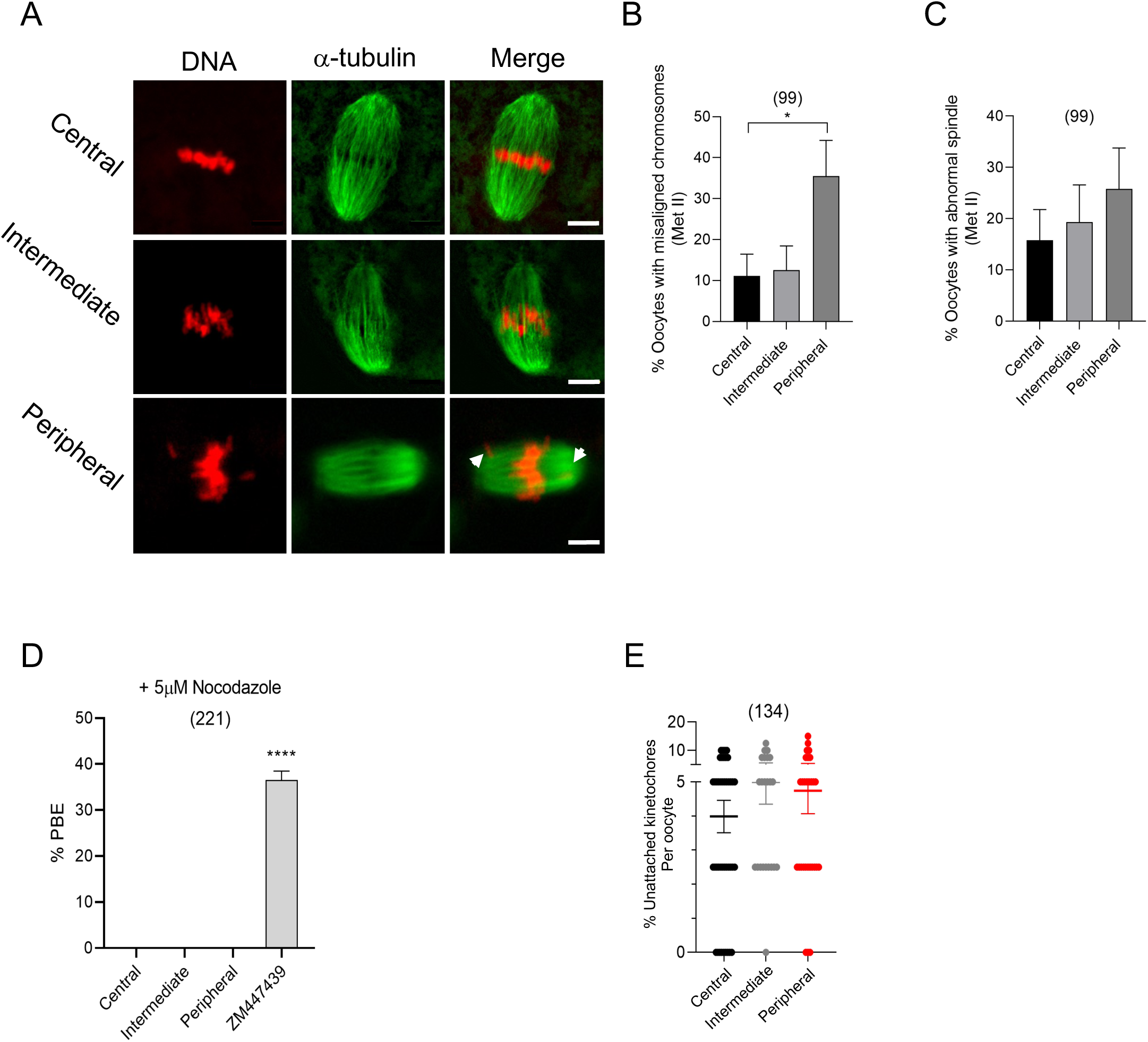
Full-grown prophase I oocytes were collected and sorted into central, intermediate and peripheral groups based on nucleus positioning and in vitro matured for 16 h (metaphase II). Oocytes were fixed and immunostained with a-tubulin to label the microtubules (the spindle). DNA was labeled by DAPI. (A) Representative confocal images of metaphase II oocytes. (B) Quantification of the percentage of chromosome misalignment in metaphase II oocytes. (C) Quantification of the percentage of abnormal spindle morphology in metaphase II oocytes. Spindle morphology was assessed using the criterion that a normal meiotic spindle is barrel shaped with two clearly defined spindle poles. The scale bar represents 10 µm. (D) Quantification of successful polar body extrusion (1^st^) in oocytes treated with nocodazole. ZM447439 oocytes served as a control. (E) Quantification of the percentage of unattached kinetochores in central, intermediate, peripheral oocytes from Fig. 2A. One-way ANOVA and Tukey’s post hoc tests were performed to determine statistical significance among groups. Data are displayed as mean ± SEM. Values with asterisks vary significantly, * P <0.05, **** P <0.0001. The total number of analyzed oocytes (from at least 3 independent replicates) is specified above each graph.

**Supplementary Figure 3:**
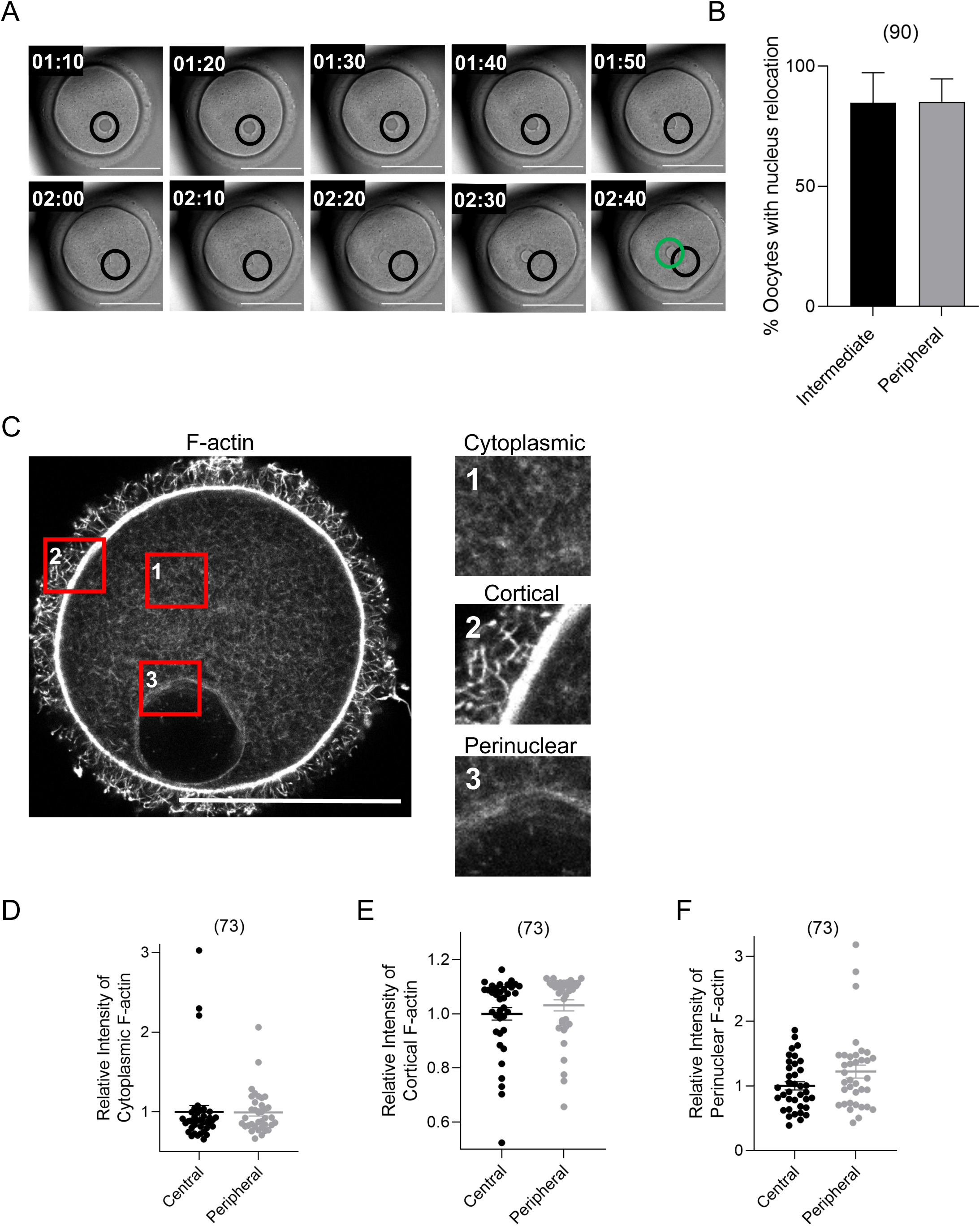
(A) Representative bright-field images of time-lapse microscopy of the movement of a peripherally positioned nucleus during prophase I arrest in oocytes incubated with milrinone. Images were taken every 10 minutes using a Z-interval of 5 µm. The black circle identifies the initial positioning of the nucleus, while the green circle denotes the final positioning of the nucleus. The scale bar represents 50 µm. (B) Quantification of the percentage of peripheral and intermediate GV oocytes achieving full relocation of the nucleus to a central position within 3 h of incubation with milrinone. (C) Representative fluorescent image of F-actin populations within oocytes. Enlarged panels featured to the right show, from top to bottom, areas containing cytoplasmic, cortical, and perinuclear F-actin. The scale bar represents 50 µm. (D-F) Quantification of the relative intensity of cytoplasmic, cortical, and perinuclear F-actin in oocytes with either a centrally or peripherally positioned nucleus. Student t-test was performed to determine statistical significance. Data are displayed as mean ± SEM.

**Supplementary Figure 4:**
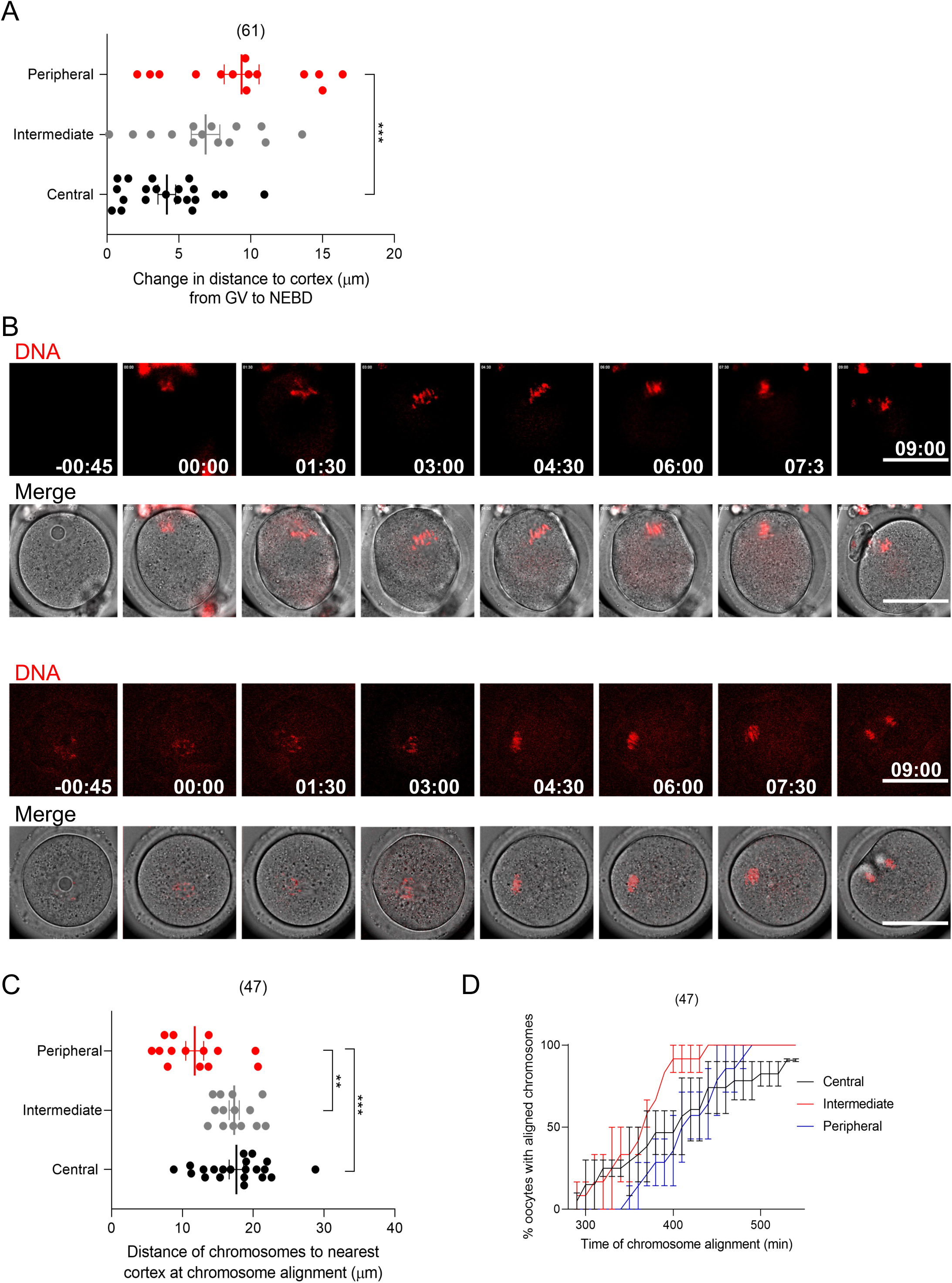
Full-grown prophase I-arrested oocytes were collected and sorted into central, intermediate, and peripheral groups based on the positioning of their nucleus and imaged live using time-lapse microscopy during *in vitro* maturation. (A) Quantification of the change in the distance of the nucleus to the nearest cortex from the initial position of the nucleus at the prophase I-arrested stage to the final destination of the nucleus immediately prior to NEBD. (B) Representative fluorescence and bright-filed image sequence depicting chromosome alignment (labeled by SiR-DNA, red) in oocytes with a peripheral (upper) and central (lower) nucleus. Images were taken every 45 minutes using a Z-interval of 7 µm. Scale bars represent 50 μm. (C) Quantification of the distance of chromosomes at the time of alignment to the nearest cortex. (D) Quantification of the time of chromosome alignment. One-way ANOVA and Tukey’s post hoc test were performed to determine statistical significance. Data are displayed as mean ± SEM. Values with asterisks vary significantly, ** P <0.01, *** P <0.001. The total number of analyzed oocytes (from 3 independent replicates) is specified above each graph.

**Supplementary Figure 5:**
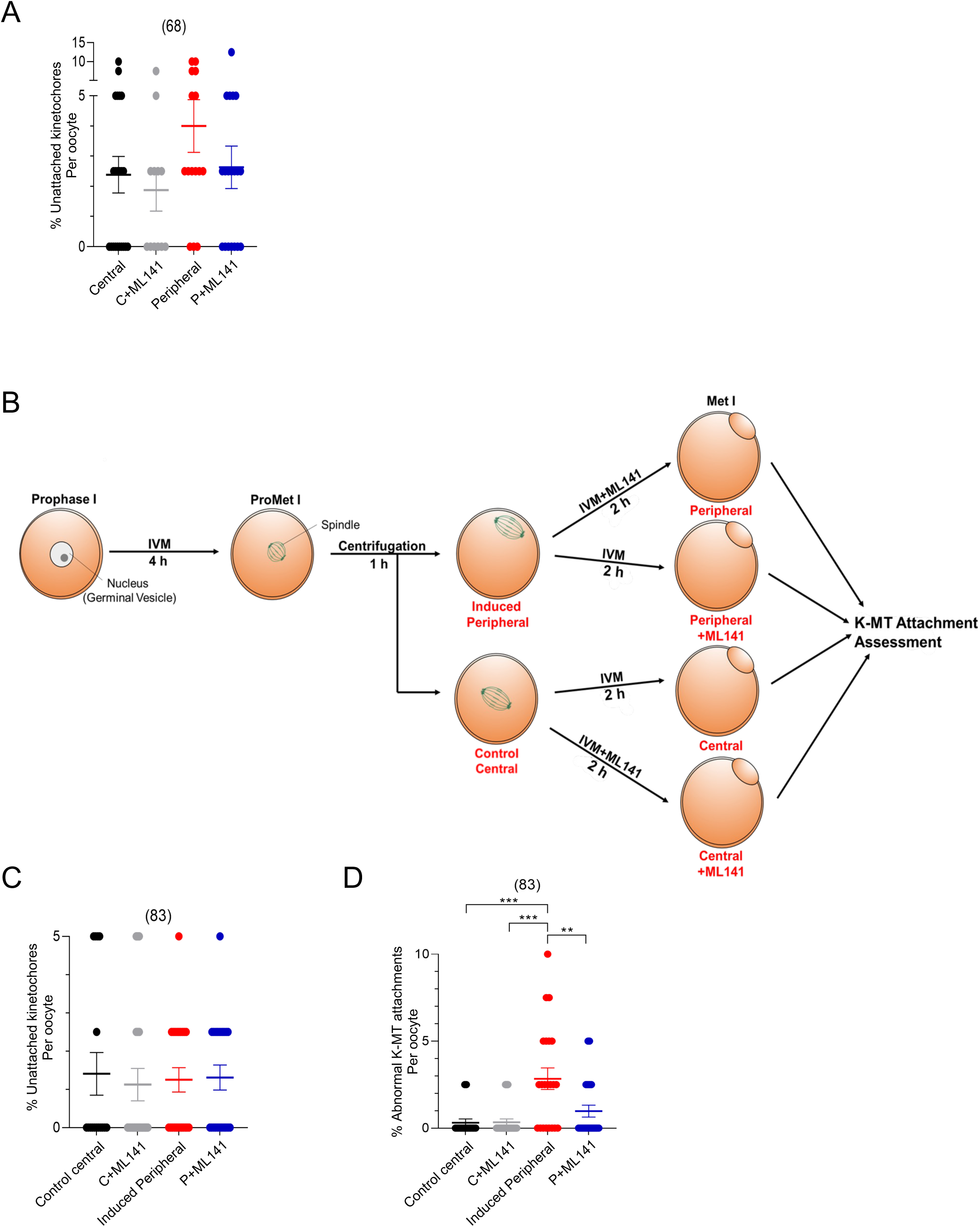
(A) Quantification of the percentage of unattached kinetochores in oocytes treated with ML141 from Fig. 3B. (B) Schematic depicting the induction of peripheral spindle positioning and treatment with ML141 in oocytes with a centrally positioned nucleus. (C) Quantification of the percentage of unattached kinetochores. (D) Quantification of the average incidence of abnormal K-MT attachment. One-way ANOVA and Tukey’s post hoc test were performed to determine statistical significance. Data are displayed as mean ± SEM. Values with asterisks vary significantly, ** P <0.01, *** P <0.001. The total number of analyzed oocytes (from 3 independent replicates) is specified above each graph.

**Supplementary Figure 6:**
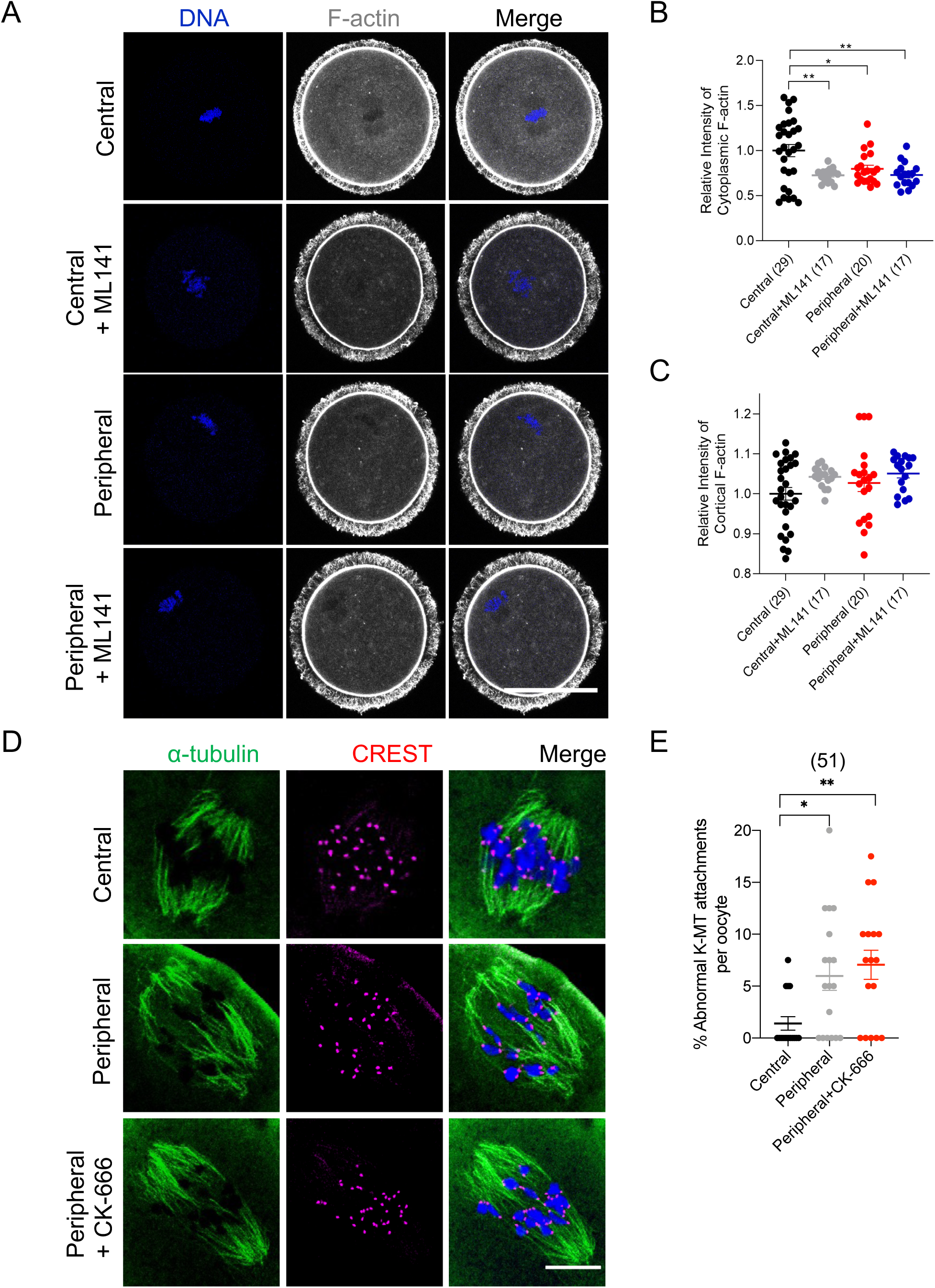
(A) Representative images of central and peripheral oocytes treated with or without ML141. The scale bar represents 50 µm. (B,C) Quantification of the relative intensity of F-actin in the cytoplasm and cortex of oocytes, respectively. (D) Representative images of K-MT attachment in prometaphase I oocytes treated with or without ARP2/3 inhibitor, CK-666 at 5 h post *in vitro* maturation. The scale bar represents 10 µm. (E) Quantification of the average incidence of abnormal K-MT attachment in D. One-way ANOVA and Tukey’s post hoc test were performed to determine statistical significance. Data are displayed as mean ± SEM. Values with asterisks vary significantly, * P<0.05, ** P<0.01. The total number of analyzed oocytes (from 4 independent replicates) is specified above the graph.

**Supplementary Figure 7:**
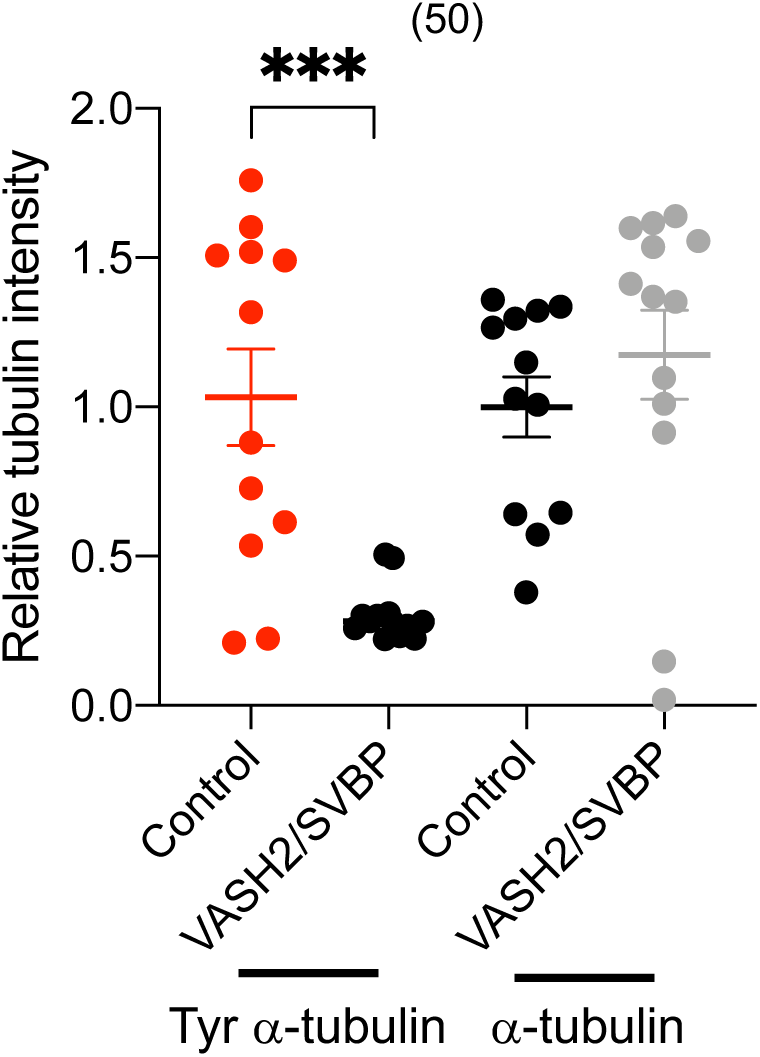
Quantification of tyrosinated α-tubulin and α-tubulin intensities in oocytes expressing VASH2/SVBP. One-way ANOVA and Tukey’s post hoc test were performed to determine statistical significance. Data are displayed as mean ± SEM. Values with asterisks vary significantly, *** P <0.001. The total number of analyzed oocytes (from 2 independent replicates) is specified above the graph.

**Supplementary Movie 1:** Time-lapse microscopy of germinal vesicle (GV) oocytes cultured in milrinone-containing medium to maintain prophase I arrest. DIC images were captured every 10 minutes at Z-intervals of 5 µm. The scale bar represents 50 µm.

## Notes

### Competing Interest Statement

The authors have declared no competing interest.

### Summary of Updates

Figure 4 is added.

